# Executive function and underlying brain network distinctions for callous-unemotional traits and conduct problems in adolescents

**DOI:** 10.1101/2023.10.31.565009

**Authors:** Drew E. Winters, Jules R Dugré, Joseph T. Sakai, R. McKell Carter

**Author notes:** Corresponding author: Drew E. Winters. Contributions Dr. Winters conceptualized the study, managed data, ran analyses, and wrote the first draft. Dr. Dugre reviewed the manuscript, provided additional citations, and revisions. Drs. Sakai and Carter both contributed to the initial conceptualization and editing revisions of the manuscript.

## Abstract

The complexity of executive function (EF) impairments in youth antisocial phenotypes of callous-unemotional (CU) traits and conduct problems (CP) challenge identifying phenotypic specific EF deficits. We can redress these challenges by (1) accounting for EF measurement error and (2) testing distinct functional brain properties accounting for differences in EF. Thus, we employed a latent modeling approach for EFs (inhibition, shifting, fluency, common EF) and extracted connection density from matching contemporary EF brain models with a sample of 112 adolescents (ages 13-17, 42% female). Path analysis indicated CU traits associated with lower inhibition. Inhibition network density positively associated with inhibition, but this association was strengthened by CU and attenuated by CP. Common EF associated with three-way interactions between density*CP by CU for the inhibition and shifting networks. This suggests those higher in CU require their brain to work harder for lower inhibition, whereas those higher in CP have difficulty engaging inhibitory brain responses. Additionally, those with CP interacting with CU show distinct brain patterns for a more general EF capacity. Importantly, modeling cross-network connection density in contemporary EF models to test EF involvement in core impairments in CU and CP may accelerate our understanding of EF in these phenotypes.

## 1. Introduction

Youth antisocial phenotypes comprise two primary components: conduct problems (CP; i.e., externalizing symptoms involving rule breaking and aggression; related to conduct disorder and oppositional defiant disorder) and callous-unemotional (CU) traits (i.e., low prosocial emotions of empathy, guilt, and remorse; related to affective dimension of adult psychopathy; for review: Blair et al., 2014). These phenotypes are often comorbid and associated with aggressive and criminal behavior, but what differentiates youth with CU are profound socio-affective impairments (Blair et al., 2014; Frick et al., 2014b), which is thought to drive persistence in antisocial behavior (Frick et al., 2014a). Likewise, these phenotypes demonstrate nuanced differences in executive functioning (EF) tasks (e.g., Dotterer et al., 2021; Fantozzi et al., 2022; Gluckman et al., 2016) and brain structures that support EF (e.g., Alegria et al., 2016; Tillem et al., 2023; Winters, Leopold, Carter, et al., 2023; Winters et al., 2021). EFs are a set of higher-order cognitive abilities that support goal-directed behavior (Smolker et al., 2015) to which impairments are transdiagnostic (Roye et al., 2022) and relate to violent criminal behavior (Meijers et al., 2017). However, EFs relationship with CU and CP remain unclear. One potential reason for this is that single EF tasks are prone to measurement error (Burgess & Rabbitt, 1997; Miyake & Friedman, 2012; Phillips & Rabbit, 1997), which, fortunately, can be redressed by modeling latent factors across EF tasks (e.g., Miyake & Friedman, 2012; Miyake et al., 2000). Another reason may be that seemingly equivalent behavioral outputs may result from brain differences and phenotypic distinctions in cognition can be revealed by examining ties to brain function (Gilman et al., 2015), specifically in contemporary EF brain models that accentuate cross-network communication (for review: O’Reilly, 2010). Thus, the present study tests how the presence of CU and CP changes associations between contemporary EF brain models and latent components of EF.

EF is comprised of multiple cognitive processes but many support a composition of three factors comprising inhibition, shifting, and fluency (Karr et al., 2019; Latzman & Markon, 2010; Roye et al., 2022; Roye et al., 2020). The healthy function of these EFs is critical for mental health (Friedman & Robbins, 2022), emotion regulation (Gyurak et al., 2012), and socio-emotional competence (Riggs et al., 2006). The impairment in these EFs, particularly top-down cognitive control such as inhibition, are ubiquitous in mental health conditions (Friedman & Robbins, 2022). It is known that EF deficits in top-down control increase potential for mental health disorders; and, despite the ubiquity, it is plausible that mental health symptoms may be characterized by distinct EF components and unique ways the brain is used to complete EF tasks (Friedman & Robbins, 2022). Thus, differences may be revealed via EF components and brain interactions underlying CU and CP.

CU and CP’s association with EF and associated functional brain properties in youth is complex. For example, when examining individual tasks, Dotterer et al. (2021) revealed neither CU traits or CP in youth associated with performance on the no/no-go or stop signal tasks, but did indicate an interaction between CU and CP such that CP associated with higher reaction times at lower CU traits and lower reaction times as higher CU traits. The authors conclude that CP uniquely contributes to sustained attention deficits during inhibitory processing tasks. Initial neuroimaging evidence by Tillem et al. (2023) demonstrates that those low in CU and high in CP had less efficiency (proportion of short connections) within the frontoparietal network, and this was partially supported by a whole brain study that did not detect frontoparietal efficiency in relation to CU (Winters, Leopold, Sakai, et al., 2023). On the other hand, Gluckman et al. (2016) demonstrated CU traits uniquely associated with decrement in cognitive control, suggesting that it is the dynamic adaptation to implement top-down control in processes such as inhibition is a unique impairment in CU traits. Neuroimaging support indicates cross-network regions involved in conflict adaption are less efficiently integrated at higher CU traits (Winters, Leopold, Sakai, et al., 2023), and that the density (number of direct connections) between conflict adaptation regions accounts for CU traits, independent of CP, via cognitive control (Winters, Leopold, Carter, et al., 2023). Importantly, Winters and Sakai (2023) demonstrate the capacity to infer another’s affect is impacted at higher CU traits when additional demands are placed on cognitive control. These findings indicate the potential for important distinctions and interactions between CU and CP as well as brain connectivity that account for executive functioning deficits and may contribute to severity differences in antisocial behavior.

Adult studies on psychopathy have implemented latent EF modeling that demonstrates distinct associations between specific psychopathy facets affective deficits (specific CU related) and antisocial behavior (specific CP related). For example, Baskin-Sommers et al. (2015) found unique variance of broader psychopathy scores (i.e., Factor 1 and Factor 2) did not associate with a common factor of EF; however, further examination revealed the specific affective deficit facet (i.e., specific CU related; psychopathy checklist-revised facet 2 – Affective: Hare et al., 1990) accounted for deficits in common factor EF above all other facets of this measure (Baskin-Sommers et al., 2015). Similarly, Fournier et al. (2021) found EF tasks with affective stimuli that revealed that the affective deficit facet (CU specific) uniquely associated with worse inhibition and affect impairments above other antisocial facets. Friedman et al. (2021) used latent modeling and found the broader antisocial factor associated with lower common EF but they were unable to separate the affective deficit facet (see the articles supplemental). This finding may tie to other work suggesting EF deficits are more widespread in relation to CP but more specific in CU traits (Baliousis et al., 2019), which is echoed by Baskin-Sommers et al. (2022) who proposes a model positing that affective processing influences EF. It is unclear if findings in Winters and Sakai (2023) or Fournier et al. (2021) support Baskin-Sommers et al. (2022) model because we do not know if the underlying effect is due to salience detection (supporting the model) or top-down control (alternative explanation). Overall, this adult literature highlights the importance of modeling specific facets of antisocial symptoms for understanding the relationship between EF and both CU and CP.

Opportunities to improve our understanding of EF impairments are highlighted by adult meta-analysis discrepancies. For example, one meta-analyses indicates broader psychopathy factors for affective deficits (broadly CU related) have moderate to small decrements in inhibition (Gillespie et al., 2022) whereas another indicates only the broader antisocial factor (broadly CP related) associate with EF deficits (Burghart et al., 2023). There is substantial heterogeneity across studies included in these meta-analyses that limit cross-study inferences because, as Morgan and Lilienfeld (2000) state in their meta-analysis of EF in psychopathy, the mean effect does not properly represent the data. Given this, cross-validation could plausibly improve meta-analysis estimates in this line of study (e.g., Willis & Riley, 2017) by using study heterogeneity to examine effect reliability instead of imposing assumptions of normal distributions (i.e., random-effect regression). Importantly, Gillespie et al. (2022) points out many of these studies compare groups with arbitrary and varied definitions. These arbitrarily defined groups by symptom extremes (1) dichotomize continuous variables that loses explanatory information (Nuzzo, 2019), (2) impose assumptions of homogeneity that are demonstrably inaccurate (e.g., Dotterer et al., 2020; Winters, Leopold, Sakai, et al., 2023), and (3) are misrepresented by means to which additional steps are required to reliably compare statistically (e.g., Liu & Wang, 2021).

These meta-analytic discrepancies increase the potential for bias related to sample characteristics. For example, a recent meta-analysis examined facet-level effects where one of the four studies used a community sample and the other three used a forensic sample (Gillespie et al., 2022), where the three forensic sample studies did not detect and effect (Maurer et al., 2016; Steele et al., 2016; Weidacker et al., 2017) and the one community sample study did find that an affective deficit (specific CU related) uniquely contributed to inhibition deficits (Fournier et al., 2021). While this could reflect sample differences, it is plausible these results are influenced by incarcerated sample characteristics such as floor effects of neurocognitive capacity (lower IQ and variance; e.g., Andover et al., 2011), celling effects of complex mental health issues (complex trauma and confounding mental health issues: e.g., Wolff & Shi, 2012), or non-random demographic factors (socioeconomic origins: e.g., Massoglia et al., 2013) that are pronounced in those that are incarcerated. This highlights the need to leverage samples with adequate variance and modern methods to bolster inferences on EF.

EF inferences distinguishing CU and CP can be bolstered by examining brain communication patterns based on a contemporary understanding of EFs in the presence of CU and CP. Critical developmental features of adolescent brains involve communication between regions (Ernst et al., 2015; Uddin et al., 2011). Brain interactions supporting EFs are emphasized in contemporary models that cut across canonical networks (for review of models: O’Reilly, 2010), and between network interactions have been found to be important for understanding both psychopathy (Dotterer et al., 2020) and CU traits (Winters & Hyde, 2022; Winters, Leopold, Carter, et al., 2023; Winters et al., 2021). Characterizing these connections may include any number of network properties (e.g., Bassett & Bullmore, 2009); however, a metric such as efficiency is thought to be unhelpful for explaining the brain (Poldrack, 2015) and, because it describes the shortness of connections, efficiency is unlikely to reliably capture the longer connections involved in cross network models of EF. Therefore, the metric of connection density is plausibly more relevant because it describes all connections without imposing assumptions on how those connections occur. Such a brain metric can help us understand performance differences between mental health symptoms despite similar performance (Gilman et al., 2015) that could clarify the complexity in the literature. Finally, latent factor modeling of EF explains the EF domain with variance across related tasks that are prone to measurement error alone (Miyake et al., 2000). Therefore, incorporating these factors in adolescent research also has the potential to help understand discrepancies in the literature.

The present study examines connection density in modern EF brain models in relation to latent EF factors and changes in the presence of CU and CP. We hypothesize that CU and CP will have distinct associations with separate EF components (inhibition, shifting, and fluency) as well as distinct interactions. Consistent with Tillem et al. (2023) we hypothesize that a three-way interaction for connection density*CP by CU will be present for inhibition as well as a common factor for EF. Finally, consistent with assertions by Baskin-Sommers et al. (2022), we hypothesize that CP will be associated with a common EF factor (representing global EF impairments) while CU will be associated with specific EFs. This work has implications for understanding the complex relationship between the brain and EFs in CU and CP.

## 2. Methods

### 2.1. Sample

Raw data from participants between the ages of 13-17 with a WAIS-II (α = 0.96; Wechsler, 2011) IQ score of >80 in the Nathan Kline Institute’s Rockland study was downloaded from the 1000 connectomes project (www.nitrc.org/projects/fcon_1000/). From a total of 122 participants, 10 were removed for IQ leaving 112 participants that were an average age of 14.52±1.31 years and predominantly White (White= 63%, Black = 24%, Asian = 9%, Indian = 1%, other= 3%) with marginally more males (female = 43%). All participants provided written consent and assent (for study procedures see: Nooner et al., 2012).

### 2.2. Measures

#### 2.2.1. Callous-Unemotional Traits

The Inventory of Callous-Unemotional traits total score was used to assess CU traits (Frick, 2004). The same factor structure validated by Kimonis et al. (2008) was used that removed two items due to poor psychometrics leaving a total of 22 items. This factor structure had adequate reliability in the current sample (Ω= 0.93).

#### 2.2.2. Conduct Problems

The externalizing subscale of the Achenbach Youth Self-Report (Achenbach and Rescorla, 2001) was used to assess CP. Validity and reliability of the externalizing measure are acceptable (Achenbach and Rescorla, 2001) and was internally consistent in the present sample (Ω= 0.75). We used the raw scores for analysis as suggested.

#### 2.2.3. Executive Function

The Delis-Kaplan Executive Function System (D-KEFS) is a battery of nine neuropsychological tests designed to assess EF (Delis et al., 2001). D-KEFS has been nationally normed for those 8-89 years old (Lezak et al., 2004) with adequate validity and reliability (Delis et al., 2004). We used the three-factor structure validated in prior work (Karr et al., 2019; Roye et al., 2022; Roye et al., 2020) including adolescents (Latzman & Markon, 2010) consisting of inhibition, shifting, and fluency. As indicated in this prior work (Karr et al., 2019; Roye et al., 2022; Roye et al., 2020), we regressed out variance for processing speed, covariation of related indicators across factors, and language function from manifest indicators and retained these residualized factors to better represent the latent executive function factor. This three-factor model had adequate reliability in the present sample (inhibit Ω= 0.79, shift Ω= 0.70, fluency Ω= 0.67). Finally, the correlation structure of the three-factor solution indicated a common factor representing a general EF capacity found in prior work (Friedman et al., 2008; Miyake et al., 2000). This common factor had adequate reliability in the present sample (Ω= 0.79). *Specific details on EF factor structure are in supplemental methods*.

#### 2.2.4. Covariates

We controlled for sex, pubertal stage, and socioeconomic status (SES). Pubertal stage and sex were measured by the Tanner assessment (Petersen et al., 1988) where parents rated pictures depicting secondary sex characteristics on a scale of 1 (pre-pubertal) to 5 (full maturity), which had adequate reliability for the present sample (Ω= 0.79). Because puberty varies by age (about five years, Parent et al., 2003) and hormonal changes during puberty impact the brain the directly impact behavior (Cameron, 2004; Dahl, 2004; Sisk and Foster, 2004), we controlled for pubertal stage instead of age. We included sex because it associates with differences in brain structure among youth with CU traits (Raschle et al., 2018). Finally, SES was assessed using the Hollingshead four-factor index (Hollingshead, 1975), which was included because of its association with EF (Lawson et al., 2018).

### 2.3. Imaging Data

#### 2.3.1. Imaging acquisition, preprocessing, and S-GIMME network estimation

All imaging data methods can be found under supplemental methods.

### 2.4. Region of Interest (ROI) selection

#### 2.4.1. Inhibition ROI’s

Contemporary understanding of inhibition involve brain models that cut across canonical networks (for review: O’Reilly, 2010). These models involve the lateral prefrontal cortices (dorsolateral and ventrolateral prefrontal cortices [dlPFC and vlPFC]) and the anterior cingulate cortex (ACC) (for review: Friedman & Robbins, 2022). For example, Banich (2019) cascade of control model proposes involvement of the dlPFC for attention task set, the vlPFC for stimulus feature representations, and the ACC for response selection and evaluation. Similarly, the expected value of control model (Shenhav et al., 2013) proposes the importance of the ACC in interaction with the dlPFC and vlPFC to allocate control to a task. Recent work on CU traits are consistent with these models indicating the ACC and cortical regions are less functionally integrated (Winters, Leopold, Sakai, et al., 2023) and may be implicated in cognitive control differences (Winters, Leopold, Carter, et al., 2023). Thus, we use these ROI’s to model inhibition. Importantly the vlPFC has anterior and mid divisions that are involved in control processes (Badre et al., 2005; Barredo et al., 2015) that are included in our ROIs.

#### 2.4.2. Shifting ROI’s

Multiple lines of research have demonstrated the medial prefrontal cortex (mPFC) and ACC to set shifting (for review: Bissonette et al., 2013). Similar to the inhibition model above, the ACC is involved in selection particularly during changing circumstances (Bissonette et al., 2013). The mPFC works in concert with the ACC to resolve set-shifting problems when attention needs to be shifted (Birrell & Brown, 2000; Bissonette et al., 2013). Therefore, these ROI’s are included to model shifting.

#### 2.4.3. Fluency ROI’s

Primary regions involved in fluency include the dlPFC and the posterior parietal cortex (PPC). The dlPFC is integral to fluency performance (Birn et al., 2010; Phelps et al., 1997) that when damaged (Baldo et al., 2001) or modulated (Ghanavati et al., 2019) impacts fluency test performance. The PPC is implicated in switching during fluency tasks (Gurd et al., 2002). Like the dlPFC, modulation of the PPC with transcranial direct current stimulation impacted fluency performance (Ghanavati et al., 2018; Ghanavati et al., 2019). Both the dlPFC and the PPC were included for fluency.

### 2.5. Analysis

All analyses were conducted in the statistical language R (Version 4.3.0; R Core Team, 2023) using the ‘lavaan’ (Rosseel, 2012) and ‘GIMME’ (Lane et al., 2021) packages. First, we extracted network features from the S-GIMME networks, then conducted diagnostics on missing data and all variables. Confirmatory factor analysis determined latent EF structure using factor loadings (λ) and model fit statistics. Factor loadings were compared to expected λ from prior work. Model fit was assessed using Hu and Bentler (1999) criteria who suggest a SRMR value < 0.08 and RMSEA < 0.06 as well as a cutoff of ≥0.95 for the CFI/TLI indicated good fit. Diagnostics of modeled variables indicated no non-linear associations or residual dependence; thus, we used maximum likelihood to estimate latent factors and regression coefficients. Confidence intervals were derived using 5000 bias-corrected bootstraps including our tests for interaction effects. Interactions were derived using the centered residual approach (Little et al., 2006; Little et al., 2007). We controlled for type I error by adjusting p values with the false discovery rate (Benjamini & Hochberg, 1995).

#### 2.5.1. Missing Data and Power Analysis

All information for power (Supplemental Figure 1) and treatment of missing data are in supplemental methods.

#### 2.5.2. Modeling Approach

We made the following modeling choices to bolster statistical inferences. ***First***, we modeled connection density as an independent variable predicting the EF outcome. This approach uses the properties of prior computationally derived brain models (i.e., model-based: Poldrack, 2011) to predict an outcome (Holdgraf et al., 2017; Naselaris et al., 2011), which allows us to infer if variance is primarily accounted for by symptoms and demographics or if there is unique variance about the brain that can account for variance in the respective executive function domain. There is substantial evidence that development of the brain and related functional properties associates with sex (Marrocco & McEwen, 2022), pubertal stage (Vijayakumar et al., 2018), and SES (Hackman et al., 2010; Moriguchi & Shinohara, 2019), as well as symptoms of CU (Winters, Leopold, Sakai, et al., 2023) and CP (Dugré & Potvin, 2023). Therefore, modeling brain properties as independent variables accounting for associated constructs strengthens inferences on brain properties association with EFs. ***Second***, we modeled multiple EF outcomes in one model that accounts for the known variance between EF domains. Conducting multiple tests on single outcomes obscures the capacity to infer whether independent variables account for unique variance on the specific EF domain because it does not account for expected shared variance between constructs. Modeling shared variance of multiple EFs allows us to state identified associations are unique to the domain of interest. ***Third***, EFs measurement error was accounted for with latent factors. Single outcome or composite measures are prone to measurement error and low reliability making it difficult to cleanly measure the EF of interest – known as the impurity problem (Burgess & Rabbitt, 1997; Denckla, 1996; Miyake & Friedman, 2012; Miyake et al., 2000; Phillips & Rabbit, 1997; Rabbitt, 1997). Latent factor modeling addresses the impurity problem (Miyake et al., 2000). ***Fourth***, we modeled density to capture the number of direct ties between nodes (Hua et al., 2022). While a multitude of important network metrics exist (e.g., Bassett & Bullmore, 2009), they are constrained by density and only characterize a chosen way information is transferred (e.g., efficiency – higher proportion of short distance connections assumed more efficient; Achard & Bullmore, 2007). Density does not impose constraints on how information transfer occurs, which is preferable when considering dynamic processes (like EF) that involve interaction across canonical networks and improves inference on behavioral associations (Gnyawali & Madhavan, 2001) in relation to EFs (Note: see supplemental analyses indicating efficiency is unrelated to EFs but correlated with density [r=0.66, p<0.001]). ***Fifth***, we modeled interactions using the centered residual approach (Little et al., 2006). This approach retains the statistical assumption that there are no shared residuals. This approach allows us to run one analysis and interpret both direct and interaction effects (Little et al., 2006; Little et al., 2007), which reduces type I error and bolsters estimate confidence. ***Sixth***, we addressed missing data. Ignoring missing data introduces bias and reduces power; and, when it is determined there is no systematic reason for missingness, addressing missing data with modern approaches improves statistical inferences (Enders, 2010; Janssen et al., 2010; Little & Rubin, 2019). ***Finally***, we modeled brain heterogeneity that is important for CU traits (e.g., Dotterer et al., 2020; Winters, Leopold, Sakai, et al., 2023). This prevents imposing the fallible assumption that brains are homogenous and allows testing if pattern-level similarities exist across heterogenous brains.

#### 2.5.3. Transparency and Openness

This study was preregistered (https://osf.io/ha5re) and code for the analysis is freely available (https://github.com/drewwint/publication_latent-EF_and_brain_in_conduct_CU_traits).

## 3. Results

### 3.1. EF Latent Factor Structure Confirmed

Confirmatory factor analysis of the three-factor EF model indicated good fit (X^2^(24) = 23.32, CFI = 1.0, TLI = 1.01, RMSEA = 0.0, SRMR = 0.068; Figure 1). Factor loadings (λ) and correlation between factors were commensurate with prior work (e.g., the tower total achievement had the lowest λ for inhibition; Karr et al., 2019). The high correlations between EF factors (r = 0.27 – 0.56) indicated the presence of a common factor found in prior work. We tested a higher order structure for the common factor (Figure 1) that resulted in the same model fit, and the common factor λ’s were commensurate with prior work (e.g., the inhibition factor loaded close to 1.0; Friedman et al., 2008; Friedman & Robbins, 2022; Miyake et al., 2000).

**Figure 1.**
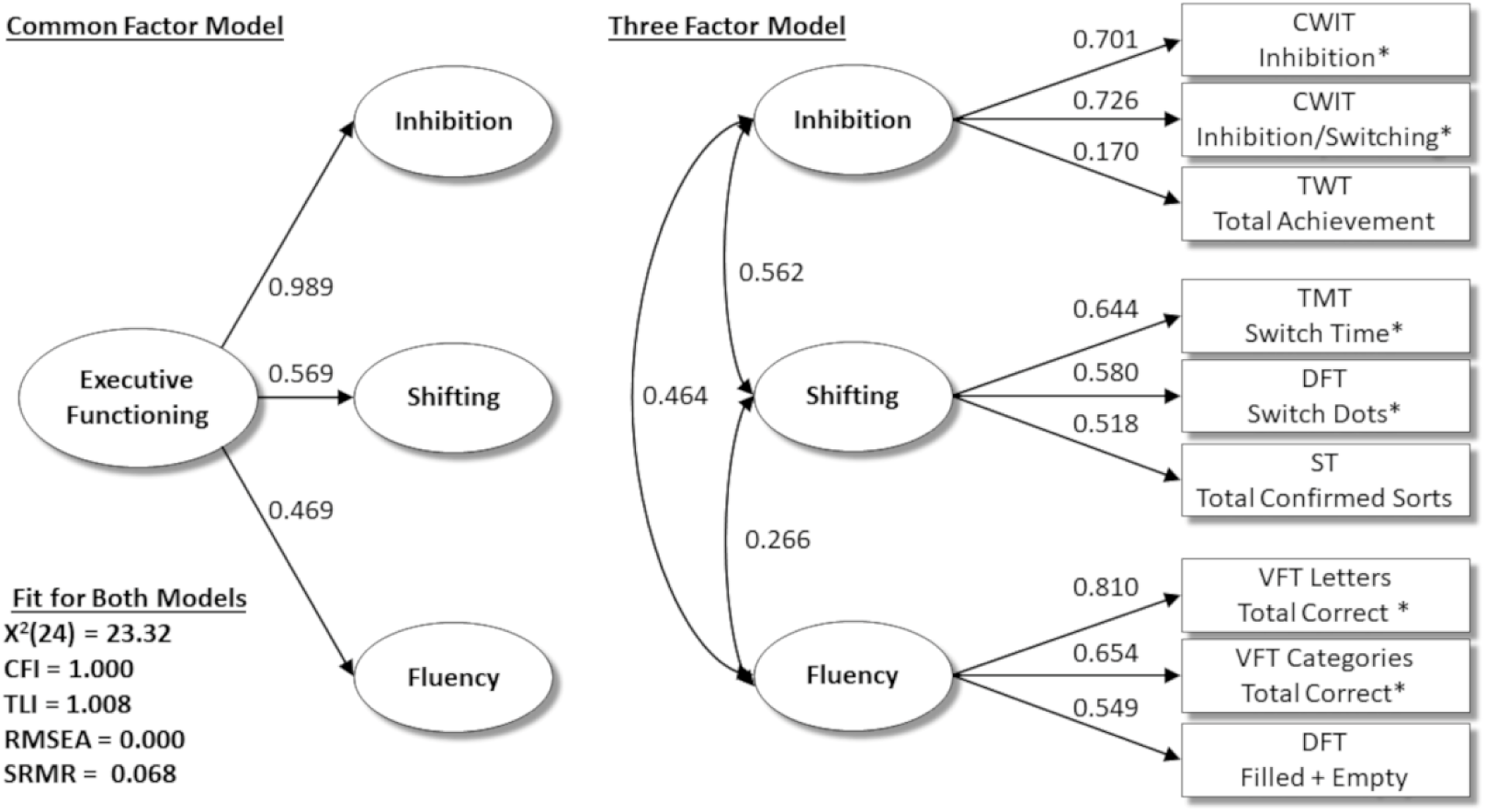
Depicting CFA of both three-factor and higher order EF model.

### 3.2. Inhibition Distinctly Associates with CU and Density Interactions with CU and CP

#### 3.2.1. Inhibition Factor

Results indicated the model accounted for 53% of the variance in inhibition (Table 1). Direct associations indicated inhibition was negatively associated with CU traits (std β = – 0.232, q= 0.044) while positively associated with both density in the inhibition network (std β = 0.287, q= 0.009) and SES (std β = 0.338, q= 0.010). Interactions indicated inhibition network density by CU (std β = 0.327, q= 0.009) and CP (std β = – 0.200, q= 0.044) were statistically meaningful (Figure 2). Network differences unique to CU traits included greater density with the left anterior vlPFC and bilateral dlPFC and right posterior vlPFC whereas connections unique to CP included less density with the bilateral dlPFC and left anterior vlPFC (Figure 3A; Supplemental Figure 2) . Simple slopes indicated the relationship between density and inhibition was highest at high CU traits and slightly attenuated at lower CU traits (Table 1; Figure 3 B) whereas for CP this association was strongest at low CP and attenuated at high CP (Table 1; Figure 3C). No three-way interactions were detected with inhibition (Table 1).

**Figure 2.**
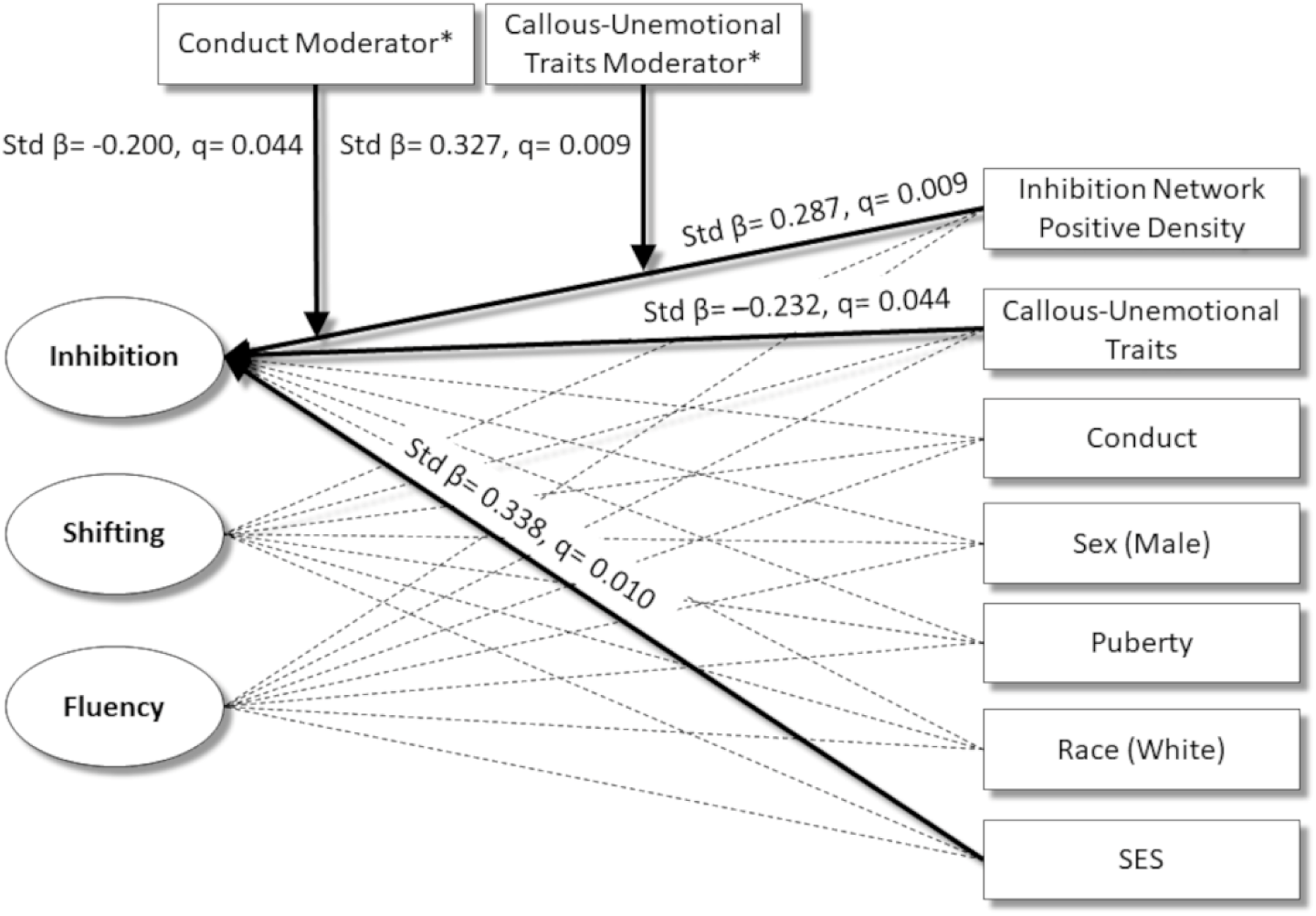
Depicting significant regressions for separate EF factors. Note *= residualized centered interaction terms; q= false discovery rate corrected significance value.

**Figure 3.**
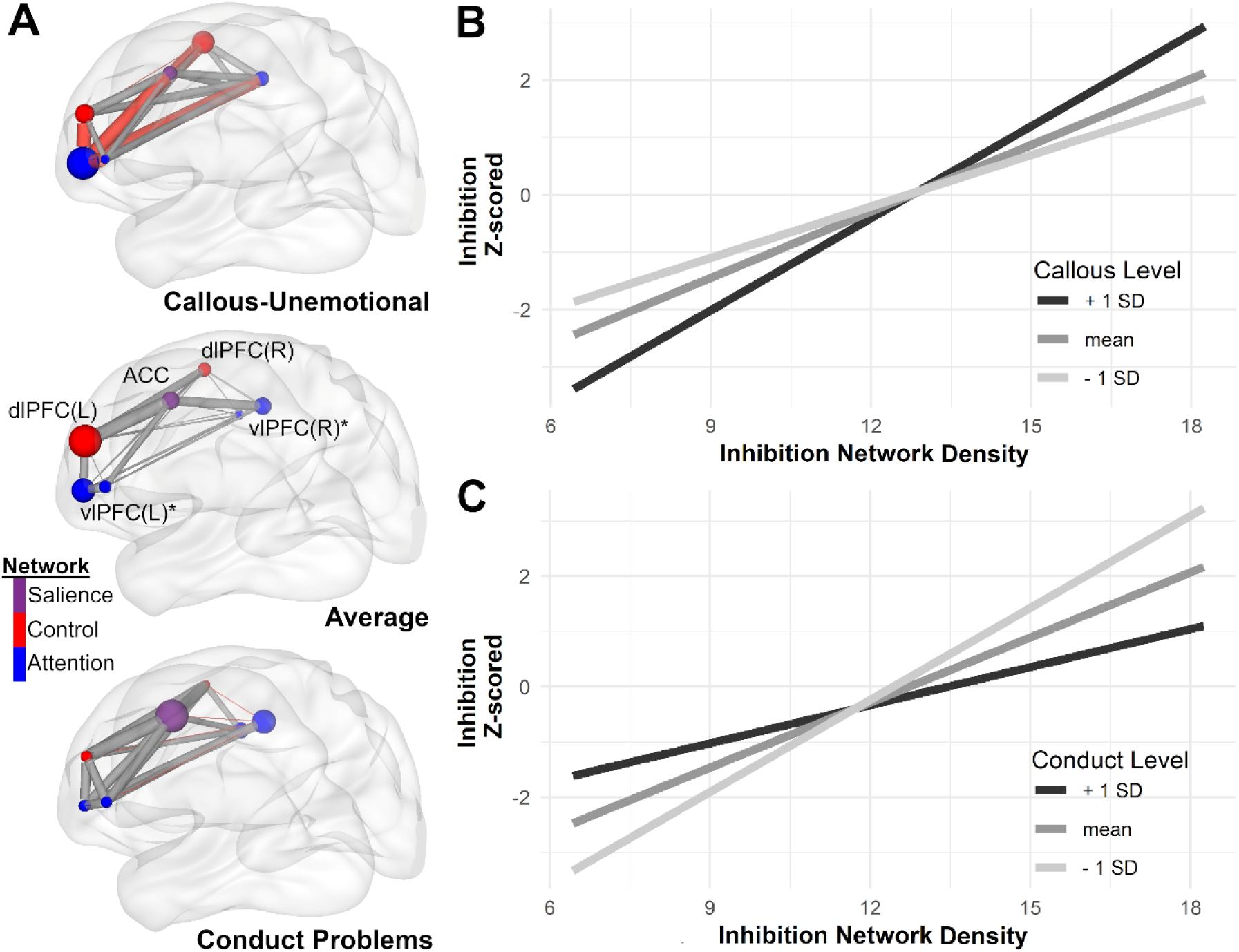
Depicting interactions between callous-unemotional (CU) traits and conduct problems (CP) with the inhibition network**. <u>(A)</u>** Depicts brain network connection density and hubness of nodes specific to CU traits (top) average for the sample (middle) and specific to conduct problems (CP) (bottom). The CU brain depicts the association of that CU with the inhibition network independent of CP and the inverse is true for the CP brain. The greater line thickness indicates greater density, and a red line indicates a significant difference that is unique to that trait. The size of each node indicates the relative importance (or hubness) of that node**. <u>(B)</u>**Depicts the interaction between CU traits and inhibition network density (independent of CP) in relation to the laten inhibition factor. **<u>(C)</u>** Depicts the interaction between CP and inhibition network density (independent of CU) in relation to the laten inhibition factor. Note: dlPFC = dorsolateral prefrontal cortex, ACC = anterior cingulate cortex, vlPFC indicates ventrolateral prefrontal cortex, (L) = left, (R) = right, and * = both anterior and posterior nodes for the vlPFC. The network depicted is the inhibition model that crosses across canonical networks and the colors for the networks indicate the canonical networks inhibition works across.

#### 3.2.2. Shifting and Fluency Factors

No statistically meaningful associations were detected after FDR correction.

### 3.3. Common EF Brain Associations Indicate Three-Way Interactions with CU and CP

The tested model accounted for 39% of the variance in the common EF factor (Supplemental Table 1). Direct associations revealed those highest in puberty for this age group had lower overall common EF (std β = – 0.257, q= 0.029). For interactions, two three-way interactions were detected for inhibition density*CP by CU (std β = – 0.168, q= 0.029; Supplemental Figure 3) and shifting density* CP by CU traits (std β = – 0.283, q= 0.006; Supplemental Figure 4). Importantly, the simple slopes for the shifting and conduct interaction by CU traits indicated it was only statistically meaningful for low conduct. One two-way interaction was detected for fluency network density*CU (std β = – 0.385, q< 0.001; Supplemental Figure 5).

## 4. Discussion

Results from adolescents in a community sample demonstrate unique associations for CU and CP with EF, which included different interactions with the brain in relation to EF. Importantly, connection density in the brain model of inhibition positively associated with latent factor inhibition after accounting for CU, CP, sex, puberty, race, and SES. For individual EF factors, inhibition was the most relevant for associations with the brain, CU, and brain interactions with both CU and CP. For the common EF factor, three-way interactions of brain density and CP by levels of CU traits were statistically meaningful. Importantly, we modeled brain heterogeneity indicated in prior work (Winters, Leopold, Sakai, et al., 2023) that revealed meaningful pattern level similarities in EF that characterize CP and CP. Overall these findings support that there are distinct and interactive effects of CU and CP in relation to brain properties that support EF as well as differences in capacity for EFs.

### 4.1. EF Factor Structure Confirmed

CFA of the latent EF model demonstrated support for both the three-factor and common-factor models in adolescents (Figure 1). This extends adult work (Karr et al., 2019; Roye et al., 2022; Roye et al., 2020) into community adolescents. This contributes to the mounting evidence suggesting this factor structure is robust and could be used across a wide variety of samples.

### 4.2. Distinct Associations of Inhibition and Brain by CU and CP

Examining EF subcomponents revealed CU traits uniquely accounted for lower inhibition after controlling for controls. This finding supports study findings that specifically examine top-down control (Gluckman et al., 2016; Winters & Sakai, 2023) as well as broader literature on the affective dimension of psychopathy (e.g., Baskin-Sommers et al., 2015). Therefore, this result supports the view that CU is related to more specific EF dysfunction (e.g., Baskin-Sommers et al., 2022). Higher connection density between regions involved in control associated with greater inhibition whereas efficiency of this network did not associate with any EF dimension (see Supplemental), which supports the importance of modeling density for cross-network brain models for specific behavior inferences.

Importantly, CU and CP had distinct and opposite interactions with connection density in relation to inhibition as well as distinct brain patterns accounting for this. The relationship between connection density and inhibition strengthened at higher CU whereas as CP attenuated this relationship. This, in part, supports assertions by Tillem et al. (2023) that brain integration related to inhibition is lower at higher CP; however, we were unable to statistically identify a three-way interaction. While it is possible that may be due to power, the null three-way interaction could indicate the differences in brain properties between CU and CP in relation to inhibition. This later assertion of unique CU and CP brain differences is supported by substantially different brain characteristics for CU and CP depicted in Figure 3A and Supplemental Figure 2. Specifically, CU traits strengthening of connection density appears to be driven by the left anterior vlPFC and bilateral dlPFC and right posterior vlPFC; whereas CP attenuating of connections density appears driven by the bilateral dlPFC and left anterior vlPFC. The differences in hub structure suggests those with CU traits rely more on the anterior vlPFC and less on both the left dlPFC and ACC whereas those with CP primarily rely on the ACC and left posterior vlPFC (Figure 3A, Supplemental Figure 2). Overall, these findings suggest substantial differences in brain properties for CU and CP in a contemporary brain model of inhibition that may help us understand nuanced differences in the literature.

Additionally, the strengthening between the brain and inhibition at higher CU traits may appear contradictory especially given that higher CU traits associated with lower inhibition. However, heightened region activation or network communication does not mean better performance but, given the association with lower inhibition, more likely indicates a brain working substantially harder at a task it is not particularly equipped to complete (e.g., Gilman et al., 2015). This was demonstrated by Khambhati et al. (2018) that found greater EF network connectivity associated positively with cognitive task performance at a low-demand but negatively at a higher-demand. Thus, higher connectivity may impede performance because of a brain that needs to work harder (revealed under higher cognitive demands) and those higher in CU traits appear to require more resources for inhibition that impacts their ability to inhibit. This view is supported by results that decrements in capacity to infer others’ emotions at higher CU traits may be due to brain differences underlying cognitive control (Winters, Leopold, Carter, et al., 2023) and greater sensitivity to demands on cognitive control (Winters & Sakai, 2023).

Together these findings suggest that youth higher in CU traits use their brains differently during top-down control and that this impacts their capacity to engage in other tasks where top-down control is required. While top-down control is ubiquitous across a number of processes (Friedman & Robbins, 2022), inhibition deficits related to how those higher in CU traits use their brain could account for a number of core impairments specific to CU such as deficient affect processing (Winters & Sakai, 2023) and prosocial behavior (Winters, Pettine, et al., 2023). Therefore, modeling cross-network inhibition brain models may reveal how differences account for impairments defining CU traits. Given a critical need for this population is to identify specific mechanisms (e.g., White et al., 2022), such work could lead to improved treatment outcomes.

### 4.3. General EF and Interactions with CP by level of CU

The EF common factor did not directly associate with CP; however, the three-way interactions between connection density and CP at varying levels of CU was present for regions associated with inhibition and shifting. Density of the inhibition network positively associated with common EF at low CP and either high or low CU that was attenuated as CP increased. Density in the shifting network showed a different pattern such that the positive association between density and common EF was only statistically meaningful at low CP at all levels of CU. The null direct associations stand in contrast to adult findings (e.g., Baskin-Sommers et al., 2015), but may indicate an association specific to youth or these direct effects to common EF may attenuate when accounting for underlying connection density or other factors (e.g., SES) included in the present model. The brain interaction findings continue to support Tillem et al. (2023) in regard to the interaction between the brain and CP is different at different levels of CU traits but extends their work to test this effect directly with EF. In the context of prior individual factor result, this finding supports that the interaction between density and CU in relation to EF remains positive after accounting for CP, which could indicate a brain working harder to accomplish EF tasks. Overall, these results indicate the brain communicates differently in relation to CU and CP symptoms for general EF, but this did not impact common EF directly. This adds additional evidence that brain properties are a critical feature showing differences for CU and CP even if the behavioral outcome equivalent.

### 4.4. Limitations

The current results have the following limitations. First, this study can only characterize contemporaneous differences in brain properties for CU and CP when this is likely to change over time. While this project addresses an important gap with innovative methods to address issues in this literature at present, future work would improve this work by examining differences over time. Second, the present study had a modest sample size (n = 112) which was adequate for estimation of the SEM models but may have missed some interaction effects. The FDR correction may have contributed to this but reducing type I errors increased confidence in significant estimates. Future studies could recruit larger and more diverse samples. Third, we used a community sample that, although substantial work suggests community samples with CU have the same neurocognitive impairments as incarcerated samples (Viding & McCrory, 2012), may not reflect the profound impairments observed in incarcerated samples. Overall, the present work adds an important contribution for understanding EF and brain distinctions between CU and CP.

### 5.5. Conclusions

The present study demonstrates distinct CU and CP associations with the brain and EFs. Specifically, CU is directly associated with decrements in inhibition whereas CP did not. The presence of CU strengthened the relationship between inhibition network connection density with inhibition; but the presence of CP attenuated the association. The three-way interaction of connection density with CP by CU in relation to common EF highlights an important interaction and differences of these phenotypes with the brain and EF. Results suggest that those higher in CU may require their brain to work harder for worse inhibition; whereas those higher in CP may fail to recruit the brain in ways that are necessary to complete EF tasks and account for findings in poor EF performance more broadly. Future work could improve our understanding of EF in CU and CP by modeling cross-network EF brain models in relation to defining deficits requiring EF such as affect processing in CU and impulsivity in CP to understand how and where brain differences are implicated in observed deficits.

## Acknowledgements

The authors express appreciation to Dr. Naomi Friedman for comments on earlier versions of this manuscript as well as for financial support by NIMH and NIDA

