## Supplementary for "Executive function and underlying brain network distinctions for callous-unemotional traits and conduct problems in adolescents"

### Supplementary Methods

#### Imaging data

**Imaging acquisition**. Instructions for participants during resting state scanning were to keep their eyes closed without falling asleep. Images were collected with a Siemens TimTrio 3T scanner using a blood oxygen level dependent (BOLD) contrast with an interleaved multiband echo planar imaging (EPI) sequence, which included a functional resting state scan (260 EPI volumes; repetition time (TR) 1400ms; echo time (TE) 30ms; flip angle 65$^{o}$; 64 slices, Field of view (FOV) = 224mm, voxel size 2mm isotropic, duration = 10 minutes) and a magnetization prepared rapid gradient echo (MPRAGE) anatomical image (TR= 1900ms, flip angle 9$^{o}$, 176 slices, FOV= 250mm, voxel size= 1mm isotropic). No scans were removed for T1 stabilization because the Siemens sequence collections images after saturation is received.

**Imaging Preprocessing**. With the raw data, we used the CONN toolbox standard preprocessing pipeline (version 18b; Whitfield-Gabrieli & Nieto-Castanon, 2012) that uses Statistical Parametric Mapping (SPM version 12; Penny et al., 2011). The Artifact Detection Tools (ART; http://www.nitrc.org/projects/artifact_detect) flagged motion outliers for correction if framewise displacement > 0.5mm and used spike regression to control for motion outliers. Because a fast multiband sequence was used to collect images, slice timing correction was not applied (Glasser et al., 2013; Wu et al., 2011). Physiologic CSF and white matter noise was regressed out of the BOLD signal using anatomic component-based noise correction method (aCompCor; Whitfield-Gabrieli & Nieto-Castanon, 2012). MPRAGE and EPI images were co-registered and normalized to an MNI template. Images were smoothed using a 6mm Gaussian kernel. Finally, to retain resting state signals, a 0.008 and 0.09Hz bandpass filter was used (Satterthwaite et al., 2013).

During preprocessing we found that 24 participants had motion > 3mm and four had >20% of invalid scans. Because this impacts the integrity of the imaging data, we did not retain the time series of these participants. This left a total of 84 participants with full imaging data and 28 participants (25%) without imaging data.

**S-GIMME**. Network maps were derived for each participant from their individual timeseries in R (Version 4.3.0; R Core Team, 2023) using the 'lavaan' (Rosseel, 2012) and 'GIMME' (Lane et al., 2021) packages. This is a data-driven sparse modeling approach that iteratively adds network connections and uses LaGrange multipliers (Sörbom, 1989) to assess model fit to retain only the statistically meaningful connections (defined as connections that improve fit for 75% of the sample). The connections retained by this sparse modeling approach minimizes spurious contemporaneous connections generated by saturated models (Gates et al., 2010). Both contemporaneous and lagged connections are modeled simultaneously for the entire sample, subgroups (data-derived groups based on functional connections), and individuals (individual-specific connections), which results in a unified structural equation model for each participant (uSEM; Gates et al., 2011). Each participants' network of connections is derived by iteratively adding connections, assessing fit of the new network, and pruning non-significant connections that may have changed with the addition of a new connection (Gates & Molenaar, 2012). This process continues until the network fits the data well according to the excellent fit criteria by Brown (2015) which requires two out of four following alternative fit criteria to be met: root mean squared error of approximation (RMSEA)≤0.05, standardized root mean residual (SRMR)≤0.05, comparative fit index (CFI)≥0.95, or non-normed fit index (NNFI)≥0.95. S-GIMME identifies shared connectivity patterns while accounting for individual heterogeneity using the community detection algorithm walktrap (Beltz & Gates, 2017), which simulations demonstrate to be a reliable method of detecting subgroups of network patterns (Gates et al., 2017; Pons & Latapy, 2005) that consistently outperforms other algorithms (Gates et al., 2016). Moreover, community detection provides the model with more known priors that improves the search for individual connections (Beltz & Gates, 2017); thus, leveraging both individual and subgroup level network features increase reliability of network connections in comparison to other network approaches (Gates et al., 2017; Gates & Molenaar, 2012; Smith et al., 2011).

**Missing Data Analysis**. Prior to analysis, we assessed data missingness using the Visualization and Imputation of Missing Values 'VIM' package (Kowarik & Templ, 2016). We tested for Missing Completely at Random (MCAR) with the 'MissMech' package in R (Mortaza et al., 2014), which uses a method by Jamshidian and Jalal (2010) that demonstrates reliability in smaller samples. Finally, to further assess missingness, we conducted t tests between those with and without missing data on variables of interest and demographic variables.

Most participants (75%) had no missing data, However, 25% of participants were missing brain data (n=28) and only 2.6% were missing executive function data (n=3). Assessment of these missing values suggested no systematic bias for missing values because (1) we could not rule out MCAR (p= 0.610) and (2) t-tests revealed no statistically meaningful difference for those with missing data. We conclude that addressing missing values would not introduce bias. Bias is substantially mitigated when using modern missing data approaches (e.g., full-information maximum likelihood; Enders & Bandalos, 2001) when compared to deletion (Enders, 2010; Janssen et al., 2010; Little & Rubin, 2019). Specifically, simulations demonstrate that full-information maximum likelihood produced unbiased estimates with over 50% of missing data when missing at random (Schafer & Graham, 2002). Thus, we used full information maximum likelihood to retain all 112 participants.

**Power analyses.** We calculated power using the package "semPower" (Moshagen & Erdfelder, 2016) at 80% power to detect an RMSEA value of 0.8 with an alpha of 0.05 that indicated we only required 51 participants to ensure the model accurately reproduced the data. We also calculated power for standardized interaction coefficients using the "pwr" (Champely et al., 2017) package for 112 participants and alpha of 0.05 that indicated we needed an association >= 0.26 for 80% power and anything lower may have missed important effects (Supplemental Figure 1).

**Latent EF Factors**

Inhibition was represented by the Tower total achievement score and Color-Work Interference tests time-to-completion (higher scores indicate worse performance) for both inhibition and inhibition/switching. Shifting was represented by the Trail Making Test Number-Letter Switching time-to-completion score, the Sorting Test confirmed correct sort score, and the Design Fluency Test switching-total correct score. Fluency was represented by the Verbal Fluency Test total correct words score for both letters and categories, and the Design Fluency Test switching-total correct score.

For regressed out variables in the latent factors, the Color-Word Interference scores were made orthogonal to word reading and color naming trials, the Trial Making Number-Letter Switching score was made orthogonal to summed performance on Number and Letter trials, the Design Fluency Switching score was made orthogonal to the Design Fluency score, and the Verbal Fluency Scores were made orthogonal to the WASI Vocabulary score.

Supplementary Table 1. Common Executive Function Model Results

| | Unstd β | SE | Std β | p | q | 95% CI Bootstrapped Lower | Upper |
|---|---|---|---|---|---|---|---|
| **Executive Function ~ ($R^2$= 0.392)** | | | | | | | |
| CU Traits | -0.014 | 0.009 | -0.101 | 0.114 | 0.171 | -0.030 | 0.003 |
| CP | 0.026 | 0.014 | 0.115 | 0.066 | 0.116 | -0.002 | 0.054 |
| Sex (Male) | 0.124 | 0.173 | 0.050 | 0.471 | 0.530 | -0.214 | 0.463 |
| Puberty | -0.257* | 0.098 | -0.200 | 0.009 | 0.029 | -0.450 | -0.064 |
| Race (White) | 0.343 | 0.190 | 0.133 | 0.071 | 0.116 | -0.029 | 0.715 |
| SES | 0.023 | 0.011 | 0.172 | 0.034 | 0.075 | 0.002 | 0.045 |
| Inhibit Network Positive Density | 0.120* | 0.042 | 0.184 | 0.004 | 0.024 | 0.038 | 0.201 |
| Moderator: CU/Density | 0.014* | 0.005 | 0.186 | 0.007 | 0.029 | 0.004 | 0.024 |
| Moderator: CP/Density | -0.020* | 0.008 | -0.166 | 0.010 | 0.029 | -0.035 | -0.005 |
| Moderator: CP/Density*CU | -0.003* | 0.001 | -0.168 | 0.011 | 0.029 | -0.005 | -0.001 |
| Shifting Network Positive Density | -0.248 | 0.217 | -0.082 | 0.252 | 0.324 | -0.672 | 0.177 |
| Moderator: CU/Density | -0.031 | 0.022 | -0.111 | 0.152 | 0.211 | -0.073 | 0.011 |
| Moderator: CP/Density | -0.009 | 0.067 | -0.010 | 0.897 | 0.897 | -0.141 | 0.123 |
| Moderator: CP/Density*CU | -0.020* | 0.006 | -0.283 | 0.001 | 0.006 | -0.031 | -0.008 |
| Fluency Network Positive Density | -0.099 | 0.158 | -0.041 | 0.530 | 0.562 | -0.408 | 0.210 |
| Moderator: CU/Density | -0.116* | 0.025 | -0.385 | 0.000 | 0.000 | -0.165 | -0.066 |
| Moderator: CP/Density | 0.054 | 0.030 | 0.121 | 0.066 | 0.116 | -0.004 | 0.112 |
| Moderator: CP/Density*CU | 0.005 | 0.006 | 0.062 | 0.409 | 0.491 | -0.007 | 0.018 |
| **Moderation Slopes** | | | | | | | |
| **Inhibition Density by Level of CP at:** | | | | | | | |
| **Low CU Traits (-1SD)** | | | | | | | |
| - 1 SD | 0.118* | 0.042 | 0.095 | 0.004 | 0.024 | 0.037 | 0.199 |
| Mean | 0.114* | 0.041 | 0.092 | 0.006 | 0.024 | 0.032 | 0.195 |
| + 1 SD | 0.109* | 0.041 | 0.088 | 0.008 | 0.024 | 0.028 | 0.190 |
| **Mean CU Traits** | | | | | | | |
| - 1 SD | 0.117* | 0.041 | 0.094 | 0.005 | 0.024 | 0.036 | 0.198 |
| Mean | 0.110* | 0.041 | 0.089 | 0.008 | 0.024 | 0.029 | 0.191 |
| + 1 SD | 0.102* | 0.041 | 0.083 | 0.013 | 0.027 | 0.021 | 0.183 |
| **High CU Traits (+1SD)** | | | | | | | |
| - 1 SD | 0.116* | 0.041 | 0.094 | 0.005 | 0.024 | 0.035 | 0.197 |
| Mean | 0.106* | 0.041 | 0.085 | 0.011 | 0.024 | 0.025 | 0.187 |
| + 1 SD | 0.096* | 0.042 | 0.077 | 0.021 | 0.035 | 0.014 | 0.177 |
| **Shifting Density by Level of CP at:** | | | | | | | |
| **Low CU Traits (-1SD)** | | | | | | | |
| - 1 SD | 0.107* | 0.041 | 0.087 | 0.009 | 0.024 | 0.026 | 0.188 |
| Mean | 0.076 | 0.042 | 0.062 | 0.068 | 0.095 | -0.006 | 0.158 |
| + 1 SD | 0.045 | 0.044 | 0.036 | 0.309 | 0.371 | -0.042 | 0.132 |
| **Mean CU Traits** | | | | | | | |
| - 1 SD | 0.099* | 0.041 | 0.080 | 0.016 | 0.028 | 0.019 | 0.180 |
| Mean | 0.049 | 0.044 | 0.039 | 0.266 | 0.342 | -0.037 | 0.135 |
| + 1 SD | -0.002 | 0.051 | -0.002 | 0.970 | 0.970 | -0.102 | 0.098 |
| **High CU Traits (+1SD)** | | | | | | | |
| - 1 SD | 0.092* | 0.041 | 0.074 | 0.026 | 0.040 | 0.011 | 0.172 |
| Mean | 0.021 | 0.047 | 0.017 | 0.650 | 0.689 | -0.071 | 0.114 |
| + 1 SD | -0.049 | 0.060 | -0.039 | 0.415 | 0.467 | -0.166 | 0.069 |
| **Fluency Density by Level of CU Traits** | | | | | | | |
| - 1 SD | -0.065 | 0.056 | -0.053 | -1.167 | 0.319 | -0.174 | 0.044 |
| Mean | -0.181* | 0.075 | -0.146 | -2.414 | 0.028 | -0.328 | -0.034 |
| + 1 SD | -0.297* | 0.097 | -0.240 | -3.062 | 0.024 | -0.487 | -0.107 |

Note: bias-corrected bootstrapped confidence intervals with 5000 resamples
q = false discovery rate corrected p-value
*= q < 0.05

### Figures

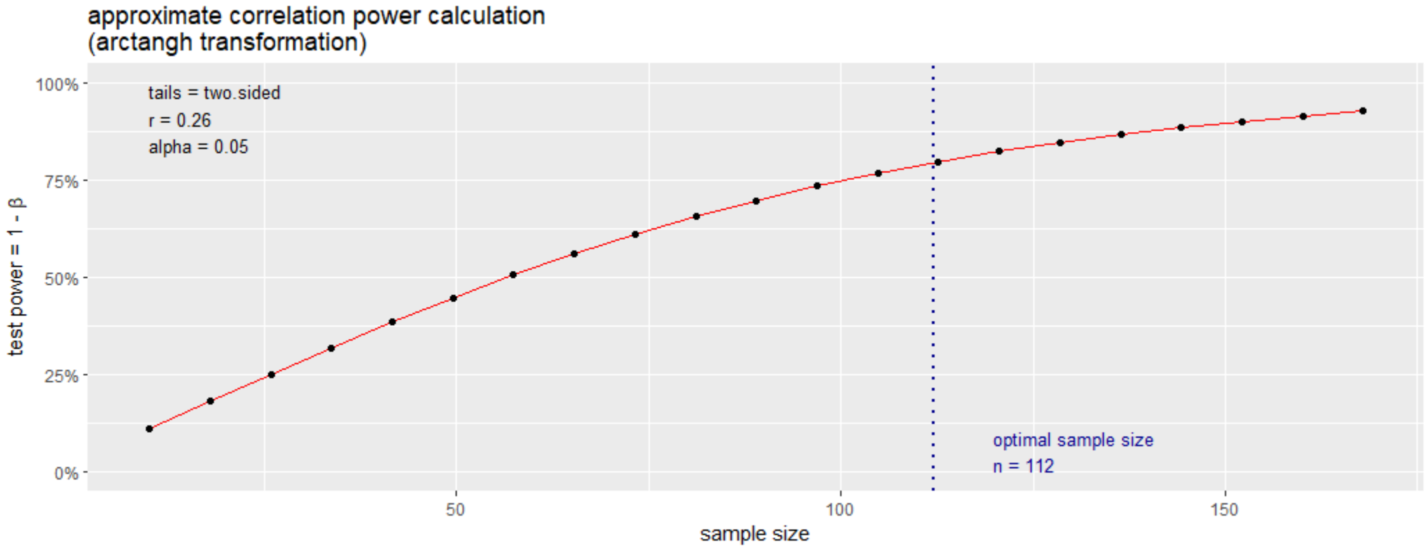


Supplemental Figure 1. Power curve for interaction effects. With the given sample size this graph indicates we require a standardized association of at least 0.26 to reach 80% power for that interaction effect.

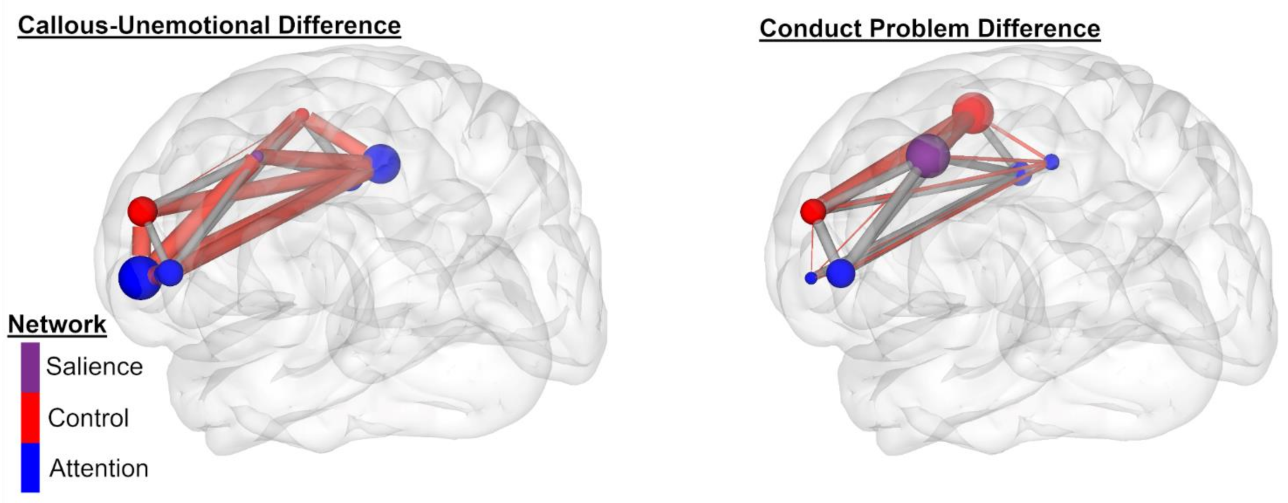


Supplemental Figure 2. Contracting differences between callous-unemotional traits and conduct problems in the inhibition network. This figure depicts the differences in one phenotype from the other. The greater line thickness indicates greater density, and a red line indicates a significant difference relative to the other phenotype. The size of each node indicates the relative importance (or hubness) of that node that relative to the other phenotype. The network depicted is the inhibition model that crosses across canonical networks and the colors for the networks indicate the canonical networks inhibition works across.

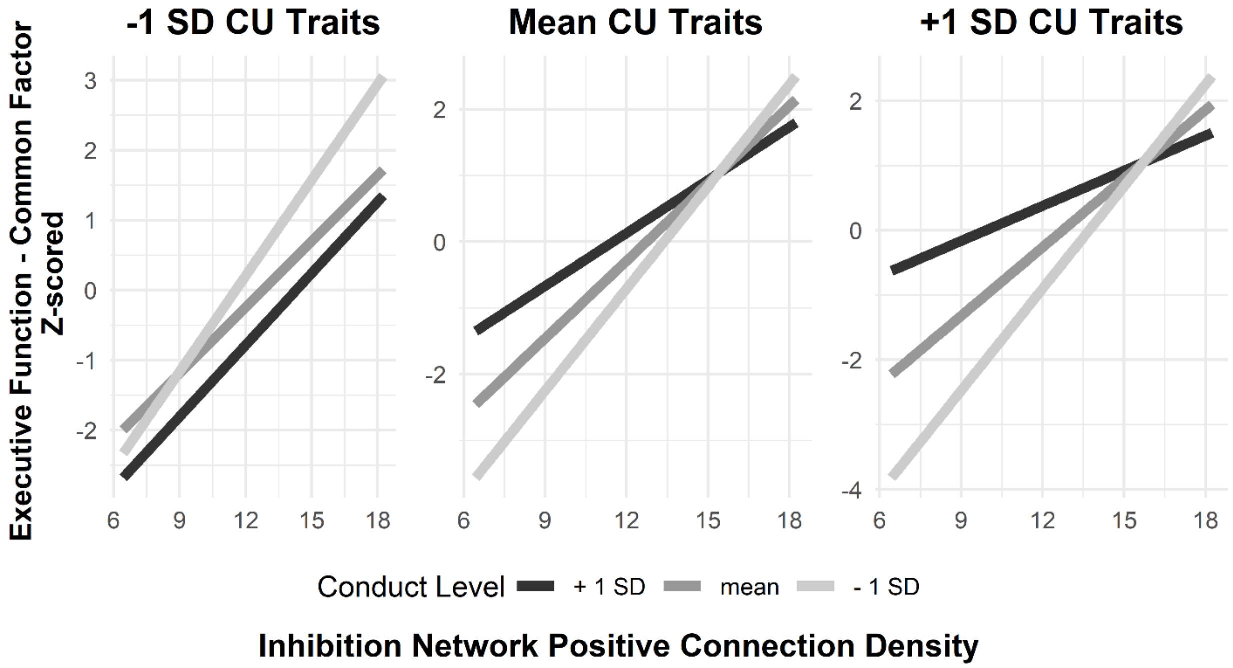


Supplemental Figure 3. Depicting three-way interaction for inhibition network density*CP by CU in relation to general EF factor.

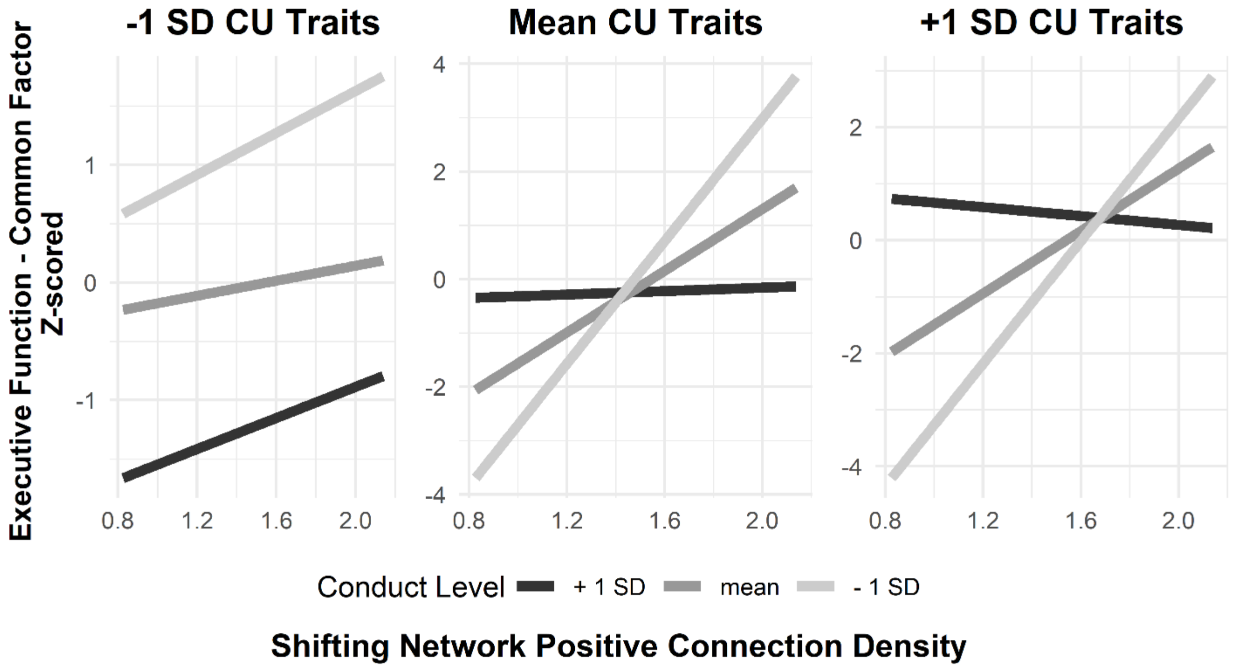


Supplemental Figure 4. Depicting three-way interaction for shifting network density*CP by CU in relation to general EF factor.

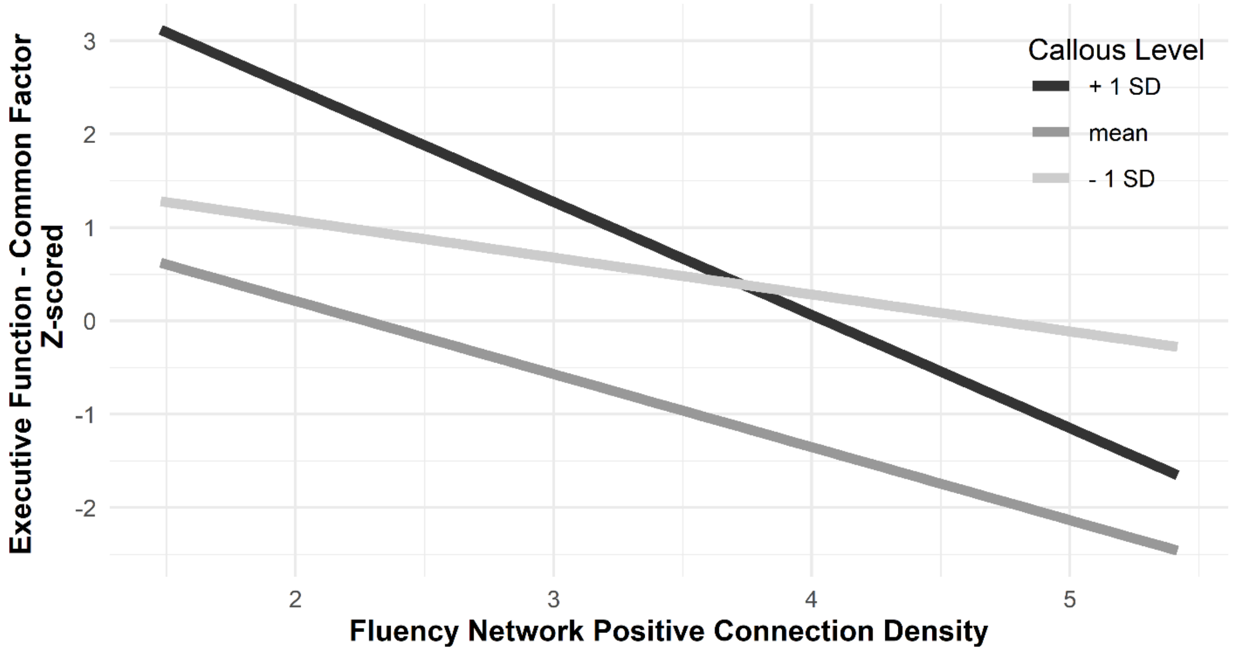


Supplemental Figure 5. Depicting interaction for fluency network density*CU in relation to general EF factor.

### Supplemental Analyses on Efficiency

Global Efficiency Analyses

| | Unstd β | SE | Std β | p | 95% CI Bootstrapped Lower | 95% CI Bootstrapped Upper |
|---|---|---|---|---|---|---|
| **Efficiency as outcome** | | | | | | |
| **Global Efficiency Inhibition Network ~** | | | | | | |
| Callous-Unemotional Traits | 0.003 | 0.014 | 0.018 | 0.855 | -0.025 | 0.030 |
| Conduct | 0.040 | 0.024 | 0.175 | 0.091 | -0.006 | 0.087 |
| Sex (Male) | -0.274 | 0.256 | -0.111 | 0.285 | -0.776 | 0.229 |
| Puberty | -0.311* | 0.138 | -0.265 | 0.024 | -0.582 | -0.040 |
| SES | 0.003 | 0.014 | 0.024 | 0.821 | -0.025 | 0.031 |
| **EF as outcome** | | | | | | |
| **Inhibition ~** | | | | | | |
| Global Efficiency Inhibition Net | 0.068 | 0.097 | 0.097 | 0.484 | -0.123 | 0.259 |
| Sex (Male) | 0.027 | 0.251 | 0.016 | 0.913 | -0.464 | 0.519 |
| Puberty | -0.038 | 0.118 | -0.042 | 0.750 | -0.269 | 0.194 |
| SES | 0.033* | 0.013 | 0.355 | 0.010 | 0.008 | 0.058 |
| **Common Executive Function ~** | | | | | | |
| Global Efficiency Inhibition Net | 0.050 | 0.062 | 0.104 | 0.417 | -0.071 | 0.172 |
| Sex (Male) | -0.144 | 0.148 | -0.121 | 0.330 | -0.434 | 0.146 |
| Puberty | 0.037 | 0.083 | 0.059 | 0.661 | -0.126 | 0.200 |
| SES | 0.038* | 0.010 | 0.583 | 0.000 | 0.017 | 0.058 |

Note: bias-corrected bootstrapped confidence intervals with 5000 resamples
*= $p < 0.05$

Local Efficiency Analyses

| | Unstd β | SE | Std β | p | 95% CI Bootstrapped Lower | 95% CI Bootstrapped Upper |
|---|---|---|---|---|---|---|
| **Local efficiency as outcome** | | | | | | |
| **Efficiency IFC anterior R ~** | | | | | | |
| Callous-Unemotional Traits | -0.013 | 0.021 | -0.065 | 0.532 | -0.054 | 0.028 |
| Conduct | -0.016 | 0.036 | -0.049 | 0.65 | -0.086 | 0.054 |
| Sex (Male) | 0.134 | 0.383 | 0.038 | 0.726 | -0.617 | 0.886 |
| Puberty | 0.108 | 0.207 | 0.065 | 0.6 | -0.297 | 0.514 |
| SES | -0.006 | 0.021 | -0.034 | 0.764 | -0.048 | 0.035 |
| **Efficiency IFC anterior L ~** | | | | | | |
| Callous-Unemotional Traits | -0.021 | 0.022 | -0.098 | 0.336 | -0.065 | 0.022 |
| Conduct | 0.073 | 0.038 | 0.206 | 0.052 | -0.001 | 0.147 |
| Sex (Male) | -0.272 | 0.406 | -0.071 | 0.502 | -1.068 | 0.523 |
| Puberty | -0.192 | 0.219 | -0.106 | 0.379 | -0.621 | 0.237 |
| SES | 0.007 | 0.022 | 0.033 | 0.762 | -0.037 | 0.05 |
| **Efficiency IFC mid L ~** | | | | | | |
| Callous-Unemotional Traits | -0.057* | 0.027 | -0.211 | 0.037 | -0.11 | -0.003 |
| Conduct | 0.038 | 0.046 | 0.087 | 0.408 | -0.052 | 0.129 |
| Sex (Male) | 0.42 | 0.498 | 0.089 | 0.399 | -0.556 | 1.396 |
| Puberty | -0.12 | 0.269 | -0.054 | 0.655 | -0.646 | 0.407 |
| SES | -0.036 | 0.027 | -0.143 | 0.192 | -0.089 | 0.018 |
| **Efficiency IFC mid R ~** | | | | | | |
| Callous-Unemotional Traits | 0.02 | 0.023 | 0.086 | 0.39 | -0.025 | 0.065 |

| | | | | | | |
|---|---|---|---|---|---|---|
| Conduct | -0.085* | 0.039 | -0.227 | 0.029 | -0.162 | -0.009 |
| Sex (Male) | 0.596 | 0.422 | 0.147 | 0.158 | -0.231 | 1.423 |
| Puberty | 0.148 | 0.228 | 0.077 | 0.514 | -0.298 | 0.595 |
| SES | 0.033 | 0.023 | 0.155 | 0.151 | -0.012 | 0.079 |
| **Efficiency dlPFC R ~** | | | | | | |
| Callous-Unemotional Traits | -0.008 | 0.017 | -0.05 | 0.634 | -0.041 | 0.025 |
| Conduct | 0.013 | 0.028 | 0.05 | 0.647 | -0.043 | 0.069 |
| Sex (Male) | 0.123 | 0.306 | 0.044 | 0.687 | -0.477 | 0.724 |
| Puberty | 0.036 | 0.165 | 0.027 | 0.826 | -0.288 | 0.36 |
| SES | -0.005 | 0.017 | -0.033 | 0.772 | -0.038 | 0.028 |
| **Efficiency dlPFC L ~** | | | | | | |
| Callous-Unemotional Traits | 0.013 | 0.023 | 0.058 | 0.578 | -0.032 | 0.057 |
| Conduct | 0.006 | 0.038 | 0.018 | 0.867 | -0.069 | 0.082 |
| Sex (Male) | 0.157 | 0.413 | 0.041 | 0.704 | -0.653 | 0.967 |
| Puberty | -0.194 | 0.223 | -0.107 | 0.385 | -0.631 | 0.243 |
| SES | 0.01 | 0.023 | 0.049 | 0.664 | -0.035 | 0.054 |
| **Efficiency ACC ~** | | | | | | |
| Callous-Unemotional Traits | -0.014 | 0.028 | -0.05 | 0.626 | -0.068 | 0.041 |
| Conduct | 0.023 | 0.047 | 0.051 | 0.63 | -0.07 | 0.115 |
| Sex (Male) | 1.027 | 0.508 | 0.216 | 0.043 | 0.032 | 2.022 |
| Puberty | 0.066 | 0.274 | 0.029 | 0.81 | -0.471 | 0.603 |
| SES | -0.004 | 0.028 | -0.017 | 0.88 | -0.059 | 0.051 |
| | | | **EF as outcome** | | | |
| **Inhibition ~** | | | | | | |
| Efficiency IFC anterior R | -0.018 | 0.034 | -0.052 | 0.601 | -0.084 | 0.049 |
| Efficiency IFC anterior L | 0.083 | 0.04 | 0.263 | 0.057 | -0.005 | 0.162 |
| Efficiency IFC mic L | -0.034 | 0.04 | -0.134 | 0.391 | -0.113 | 0.044 |
| Efficiency IFC mid R | -0.005 | 0.041 | -0.017 | 0.898 | -0.085 | 0.075 |
| Efficiency dlPFC R | -0.065 | 0.064 | -0.15 | 0.314 | -0.191 | 0.061 |
| Efficiency dlPFC L | 0.007 | 0.034 | 0.023 | 0.831 | -0.06 | 0.075 |
| Efficiency ACC | 0.019 | 0.029 | 0.077 | 0.505 | -0.038 | 0.077 |
| Sex (Male) | -0.099 | 0.126 | -0.082 | 0.433 | -0.346 | 0.148 |
| Puberty | 0.007 | 0.067 | 0.011 | 0.915 | -0.124 | 0.139 |
| SES | 0.013 | 0.013 | 0.205 | 0.311 | -0.013 | 0.039 |
| **Common EF ~** | | | | | | |
| Efficiency IFC anterior R | 0.023 | 0.031 | 0.075 | 0.468 | -0.039 | 0.085 |
| Efficiency IFC anterior L | 0.054 | 0.038 | 0.193 | 0.154 | -0.02 | 0.128 |
| Efficiency IFC mic L | 0.001 | 0.029 | 0.006 | 0.963 | -0.056 | 0.059 |
| Efficiency IFC mid R | 0.031 | 0.044 | 0.119 | 0.475 | -0.055 | 0.118 |
| Efficiency dlPFC R | -0.028 | 0.048 | -0.073 | 0.567 | -0.123 | 0.067 |
| Efficiency dlPFC L | -0.021 | 0.039 | -0.074 | 0.596 | -0.098 | 0.056 |
| Efficiency ACC | 0.024 | 0.027 | 0.109 | 0.357 | -0.028 | 0.077 |
| Sex (Male) | -0.199 | 0.132 | -0.187 | 0.133 | -0.458 | 0.061 |
| Puberty | 0.059 | 0.072 | 0.108 | 0.407 | -0.081 | 0.2 |
| SES | 0.034* | 0.01 | 0.582 | 0.001 | 0.013 | 0.054 |

Note: bias-corrected bootstrapped confidence intervals with 5000 resamples
*= p < 0.05
